# Gac is a transcriptional repressor of the Lyme disease spirochete’s OspC virulence-associated surface protein

**DOI:** 10.1101/2022.11.09.515855

**Authors:** Tatiana N. Castro-Padovani, Timothy C. Saylor, Olivia T. Husted, Andrew C. Krusenstjerna, Nerina Jusufovic, Brian Stevenson

## Abstract

The OspC outer-surface lipoprotein is essential for the Lyme disease spirochete’s initial phase of vertebrate infection. Bacteria within the midguts of unfed ticks do not express OspC, but produce high levels when ticks begin to ingest blood. Lyme disease spirochetes cease production of OspC within 1-2 weeks of vertebrate infection, and bacteria that fail to downregulate OspC are cleared by host antibodies. Thus, tight regulation of OspC levels is critical for survival of Lyme borreliae, and therefore an attractive target for development of novel treatment strategies. Previous studies determined that a DNA region 5’ of the *ospC* promoter, the *ospC* operator, is required for control of OspC production. Hypothesizing that the *ospC* operator may bind a regulatory factor, DNA affinity pulldown was performed, and identified binding by the Gac protein. Gac is encoded by the C-terminal domain of the *gyrA* open reading frame, from an internal promoter, ribosome-binding site, and initiation codon. Our analyses determined that Gac exhibits a greater affinity for *ospC* operator and promoter DNAs than for other tested borrelial sequences. In vitro and in vivo analyses demonstrated that Gac is a transcriptional repressor of *ospC*. These results constitute a substantial advance to our understanding the mechanisms by which the Lyme disease spirochete controls production of OspC.

**Importance:** *Borrelia burgdorferi* (sensu lato) requires its surface-exposed OspC protein in order to establish infection of humans and other vertebrate hosts. Bacteria that either do not produce OspC during transmission, or fail to repress OspC after infection is established, are rapidly cleared by the host. Herein, we identified a borrelial protein, Gac, that exhibits preferential affinity to the *ospC* promoter and 5’ adjacent DNA. A combination of biochemical analyses and investigations of genetically-manipulated bacteria demonstrated that Gac is a transcriptional repressor of *ospC*. This is a substantial advance toward understanding how the Lyme disease spirochete controls production of the essential OspC virulence factor, and identifies a novel target for preventative and curative therapies.

## Introduction

The Lyme disease spirochete, *Borrelia burgdorferi* sensu lato (*B. burgdorferi* sl), requires its OspC protein during the initial stages of vertebrate infection, and mutant bacteria that cannot produce this outer surface-exposed lipoprotein are unable to initiate infection of vertebrates (1-5). Production of OspC is tightly regulated. *B. burgdorferi* sl within the midguts of unfed ticks do not express OspC (6, 7). As ticks begin to feed on host blood, spirochetes initiate production of OspC, with the large majority producing detectable quantities of OspC at the time of transfer from tick to vertebrate (6, 8-11). The essential function(s) of OspC is not conclusively known - it can bind a tick salivary protein and vertebrate plasminogen and complement C4b - but may have additional functions (12-22). Infected humans and other animals mount strong antibody responses against OspC within days of infection (23-26). At the same time, *B. burgdorferi* sl represses expression of OspC, and spirochetes that do not shut off OspC synthesis are cleared by immunocompetent hosts (2, 3, 5, 27-31). Thus, regulation of OspC may be an excellent target for novel therapeutics to prevent and/or clear *B. burgdorferi* infections.

OspC was the first *B. burgdorferi* sl protein to be identified as being regulated during the natural tick-vertebrate infectious cycle (6). As noted above, OspC is not produced in unfed ticks, but is highly expressed in feeding ticks and during the earliest stages of vertebrate infection (6, 8). Regulated production can also be manipulated in culture, with bacteria grown at 23°C, pH=7.5, or culture media that yield slow replication rates, producing low to no OspC, while the protein is highly expressed following transfer from those conditions to 35°C, pH=6.5, or complete culture media, respectively (6, 32-34). It is not yet known how those culture conditions are interpreted by *B. burgdorferi* sl to modulate OspC levels, but they provide convenient means to evaluate borrelial regulatory mechanisms.

An alternative sigma factor, RpoS, is also affected by the above-described infection and culture conditions (34-37). RpoS controls production of numerous *B. burgdorferi* sl proteins that are required for vertebrate infection (36, 38, 39). Deletion of *rpoS* blocks induction of OspC, which has led to a hypothesis that expression of *ospC* is solely due to RpoS-RNA polymerase holoenzyme directly transcribing from the *ospC* promoter (36, 37, 40, 41). Yet, there is evidence that transcriptional control of *ospC* is more complicated than that model assumes. Previous studies found that a region of DNA 5’ of the *ospC* transcriptional start site, designated the *ospC* operator, is necessary for induction of *ospC* transcription (42-44).

*B. burgdorferi* sl has been divided into several clades that are often referred to as genospecies, which include *B. burgdorferi* sensu stricto (*B. burgdorferi* ss) and *B. garinii*. An earlier study noted that the *ospC* operator of *B. burgdorferi* ss type strain B31 includes a pair of inverted repeat sequences (Fig. 1) (44, 45). Yet, those inverted repeats are found in only a subset of Lyme disease borreliae (46). However, the *ospC* operators of all examined Lyme spirochetes contain two or more copies of a conserved 11bp directly-repeating sequence (Fig. 1 and (46)). *B. burgdorferi* ss B31 contains two copies of the 11bp repeat, which overlap the previously discussed inverted repeats (Fig. 1 and (44, 46)). In contrast, the *ospC* operators of some strains of the *B. garinii* genospecies contain seven adjacent copies of the 11bp repeat (Fig. 1). All *ospC* promoters also contain a copy of the 11bp repeat, oriented in the opposite direction (Fig. 1). The retention of the 11bp repeat suggested to us that it may be a key component of *ospC* regulation, with sequence conservation pointing toward it being a binding site for a regulatory protein. To address that hypothesis, we performed DNA-binding studies with borrelial cytoplasmic extract, identified a component protein that has affinity for the *ospC* operator, and identified a functional role for that protein as a transcriptional repressor of *ospC*.

**Figure 1.**
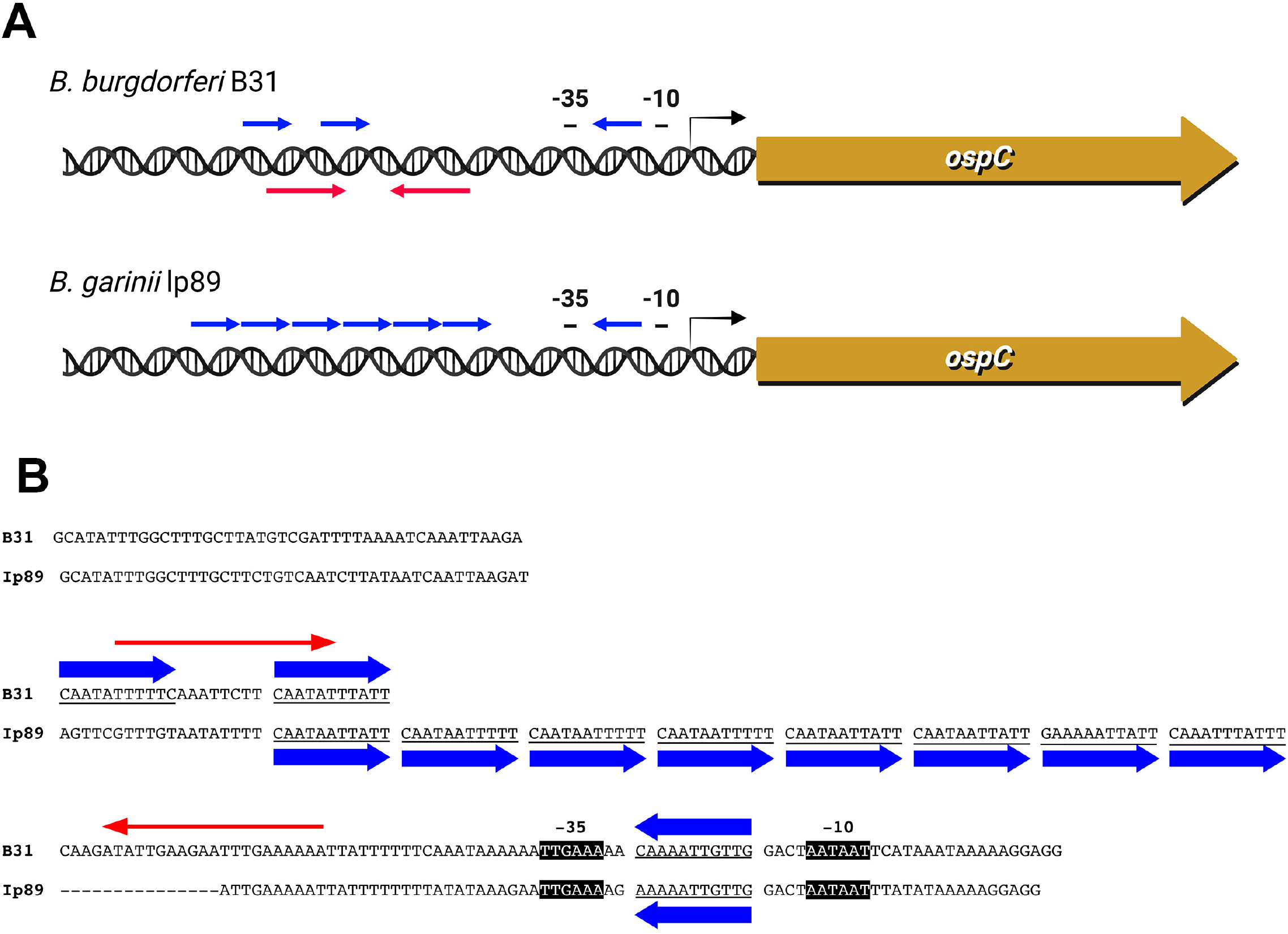
DNA sequences 5’ of the *ospC* open reading frames of *B. burgdorferi* ss B31 and *B. garinii* Ip89 contain conserved and non-conserved repeated sequences. **(A)** Graphic representation of the 11bp direct repeats (blue arrows) and a distinct, longer inverted repeat sequence in the operator of strain B31 (red arrows), that is not found in strain Ip89 or many other Lyme disease borreliae (44, 46). **(B)** Alignment of the *ospC* 5’ DNA sequences. Strain B31 contains two 11bp repeats that are separated by a unique 8bp sequence, while Ip89 carries seven directly repeated copies. In addition, both strains contain an inverted copy of the 11bp sequence between their -35 and -10 promoter elements.

## Results

### Identification of Gac as an *ospC* operator-binding protein

As a first step, an infectious clone of the *B. burgdorferi* ss type strain, B31-MI-16, was cultured to mid-exponential phase at 35°C. That condition was adopted based on our previous characterization of the *B. burgdorferi erp* operons, whose expression is also responsive to culture temperature, in which we found that the *erp* repressing and activating proteins were all adequately expressed when cultured at 35°C (47-49).

A fluorescently-tagged DNA sequence corresponding to the *ospC* operator region of *B. garinii* Ip89 was incubated with the B31-MI-16 cytoplasmic extract, then subjected to electrophoretic mobility shift electrophoresis (EMSA). The strain Ip89 sequence was used for these analyses under the hypothesis that a protein that binds to the 11bp repeated sequence would have an overall greater affinity for the 7 direct repeats of Ip89 as opposed to the smaller number of separated repeats of B31 (Fig. 1). Addition of the cytoplasmic extract resulted in retarded mobility of the labeled DNA probe, indicating that a borrelial component bound to the DNA (Fig. 2A).

**Figure 2.**
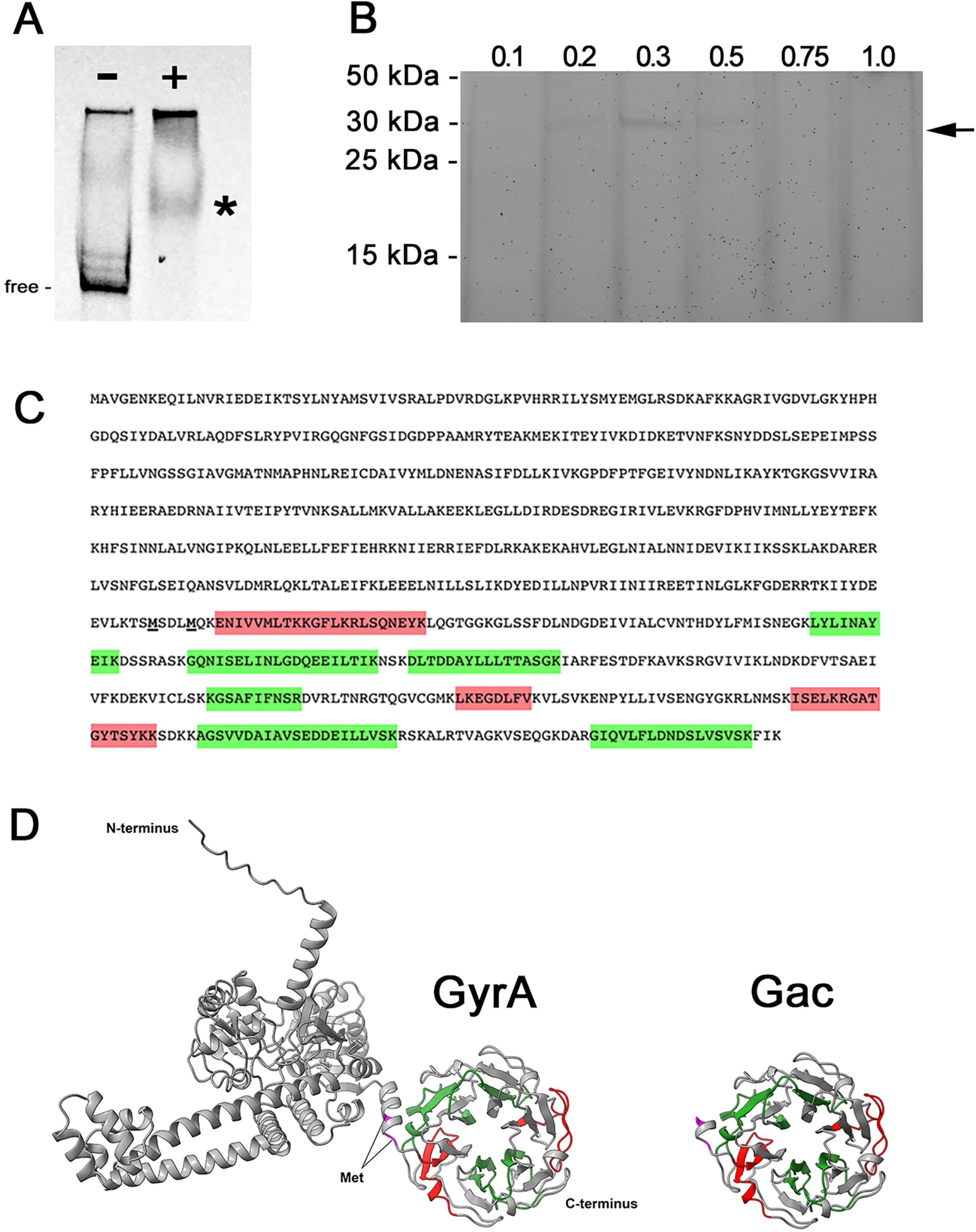
**(A)** Electrophoretic mobility shift assay (EMSA) of labeled Ip89 *ospC* operator, without (-) and with (+) cytoplasmic extract from strain B31. The asterisk indicates shifted probe that was present only when borrelial extract was included, indicating presence of a DNA-binding component. Unbound DNA is indicated by “free”. Some labeled DNA probe remained in the wells of the gel. **(B)** Cytoplasmic extract from strain B31 was incubated with Ip89 *ospC* operator DNA affixed to beads, eluted with increasing concentrations of NaCl, and eluates were separated by SDS-polyacrylamide gel electrophoresis. Lanes are labeled with the concentration of NaCl used for that elution. The arrow indicated a protein band of 30-35 kDa that eluted in the 0.3 and 0.5 mM NaCl washes. **(C)** Amino acid sequence of GyrA and Gac (the two potential initiation methionines of Gac are indicated by boldface and underlining). Mass spectrometry identified nine unique peptides that all mapped to the Gac ORF. Peptides identified with high confidence are shaded green, and those with lower confidence in red. **(D)** Predicted structures of *B. burgdorferi* Gac and GyrA proteins, as determined by AlphaFold (https://alphafold.ebi.ac.uk). Both are consistent with the solved structures of *B. burgdorferi* Gac (52) and the N-terminal domain of *E. coli* GyrA (https://www.rcsb.org/structure/4CKK) (83).

Ip89 *ospC* operator DNA was again PCR amplified, but with one of the oligonucleotide primers containing a 5’ biotin moiety. That DNA was adhered to streptavidin-coated magnetic beads, incubated with B31 cytoplasmic extract, washed extensively, then bound proteins eluted with buffers having progressively greater concentrations of NaCl. Aliquots of each eluted fraction were subjected to SDS-PAGE and proteins stained with SYPRO-Ruby. A protein band with an approximate molecular mass of 30-35 kDa eluted at 0.3 and 0.5 M NaCl (Fig. 2B). The bands were excised from the gel, subjected to trypsin digestion and mass spectrometry, and results compared to the predicted proteome of strain B31. Only one potential match was obtained, to the GyrA subunit of DNA gyrase, with 9 unique peptides identified. Notably, all of the identified peptide fragments mapped to the C-terminal half of GyrA (Fig. 2C).

Bacterial DNA gyrases are comprised of two subunits, GyrA and GyrB (50). GyrA proteins possess a unique C-terminal domain that bends DNA to facilitate the topoisomerase activity of the gyrase (51, 52). Curiously, Lyme disease borreliae produce two separate proteins from the *gyrA* locus, a full length GyrA protein and a second, 35 kDa protein named Gac, which is transcribed from an internal promoter and consists solely of the C-terminal DNA-binding domain (52-54) (Fig. 2D, and Fig. S1). We pursued the possibility that Gac preferentially binds to *ospC* operator DNA and thereby affects production of OspC.

### Gac exhibits affinity for the *ospC* operator and promoter

To address the question of Gac affinity, labeled DNA probes were constructed that consisted of either the strain Ip89 or B31 *ospC* operators (Fig. 1). Additional unlabeled DNAs were produced for use as EMSA competitors: each of the *ospC* operators, truncated segments of the B31 *ospC* operator, and DNA sequences corresponding to the 5’ ends of the borrelial *flaB* and *ospAB* operons. The Ip89 *ospC* operator probe and competitor were derived by PCR from plasmid pCR2.1 clones, using plasmid-specific M13 Reverse and Forward primers, so an additional unlabeled competitor was produced by PCR of empty pCR2.1, to account for vector-derived DNA sequences.

EMSAs of the Ip89 *ospC* operator probe and recombinant Gac found that the protein bound to the labeled probe (Fig 3A, lane 2). Densitometric analyses of EMSAs revealed that Gac exhibits differences in binding affinity to the tested sequences. Addition of 10-times excess of unlabeled Ip89 *ospC* operator resulted in a two-fold reduction in the DNA-Gac EMSA band, while 25-fold excess of unlabeled Ip89 *ospC* operator resulted in greater than an eight-fold reduction in binding and eliminated detectable levels of DNA-protein complexes (Fig. 3A, lanes 3 and 4, respectively). Addition of 25-times excess unlabeled B31 *ospC* operator had little effect on the Ip89 operator-Gac complex, indicating a greater affinity of Gac for the Ip89 operator than for the B31 operator (Fig 3A, lane 5). Note that the primary difference between the Ip89 and B31 DNAs is the number of 11bp repeat sequences in their *ospC* operators. Inclusion of 25-fold excesses of either empty pCR2.1 amplicon, *flaB*, or *ospAB* unlabeled DNAs also did not compete Gac away from the labeled probe (Fig 3A, lanes 6, 7, and 8, respectively). Thus, Gac displays greater affinity for the Ip89 *ospC* operator than it does for any of the other tested sequences.

**Figure 3.**
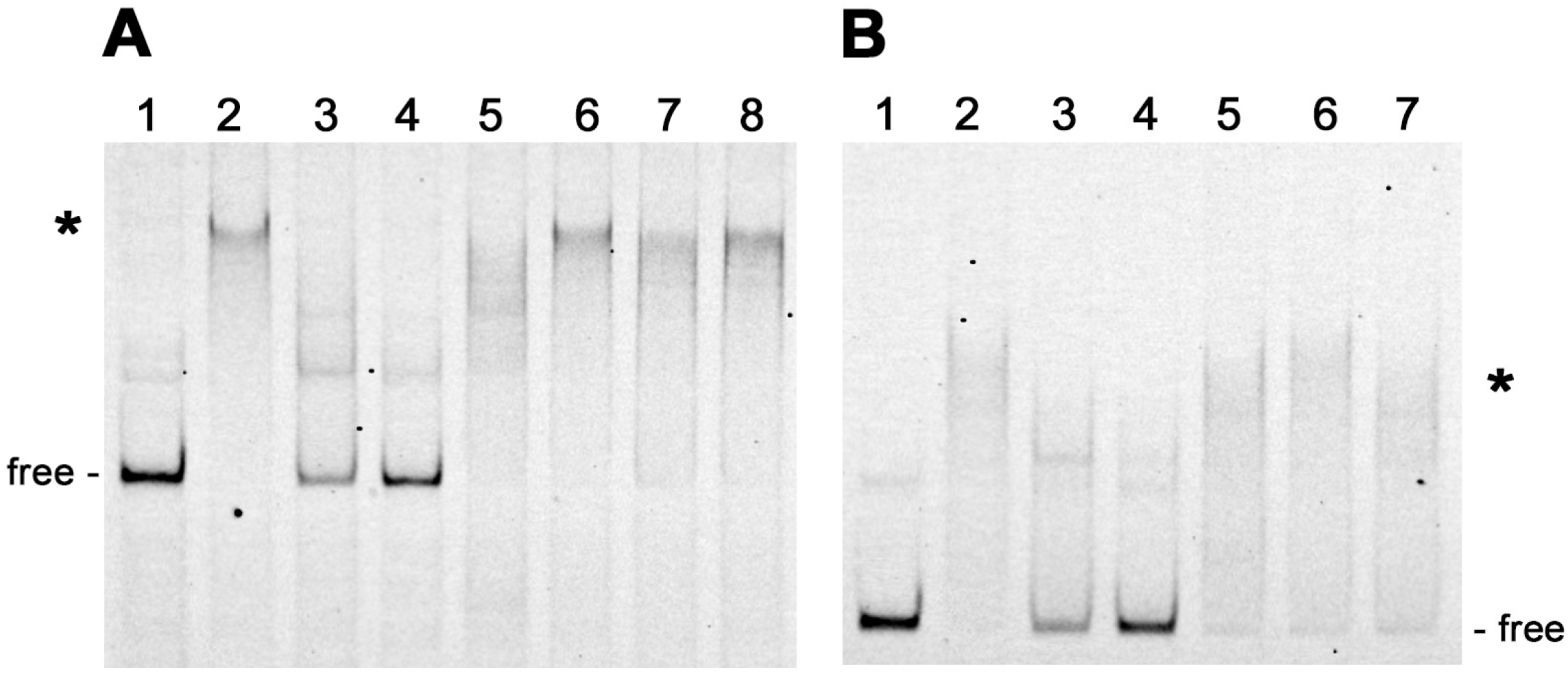
Representative EMSAs of purified recombinant Gac with labeled *ospC* operator/promoter DNAs and unlabeled competitors. Shifted DNAs are indicated with asterisks. Unoccupied DNAs are labeled “free”. **(A)** Labeled Ip89 operator/promoter (10 nM) without (lane 1) and with 25 nM Gac (lanes 2-8). Lane 2: no added competitor. Lane 3: plus 10x excess unlabeled Ip89 *ospC* operator/promoter. Lane 4: plus 25x excess unlabeled Ip89 *ospC* operator/promoter. Lane 5: plus 25x excess unlabeled B31 *ospC* operator/promoter. Lane 6: plus 25x unlabeled pCR2.1 amplicon. Lane 7: plus 25x excess unlabeled *flaB* DNA. Lane 8: plus 25x unlabeled *ospAB* DNA. **(B)** Labeled B31 operator/promoter (10 nM) without (lane 1) and with 25 nM Gac (lanes 2-7). Lane 2: no added competitor. Lane 3: plus 10x excess unlabeled B31 *ospC* operator/promoter. Lane 4: plus 25x excess unlabeled B31 *ospC* operator/promoter. Lane 5: plus 10x unlabeled pCR2.1 amplicon. Lane 6: plus 10x excess unlabeled *flaB* DNA. Lane 7: plus 10x unlabeled *ospAB* DNA.

Similar EMSAs and densitometric analyses were performed with labeled B31 *ospC* operator as probe. Unlabeled B31 *ospC* operator DNA effectively competed Gac away from the labeled probe (Fig 3B, lanes 3-4). Addition of a 10-fold excess of unlabeled B31 *ospC* operator competitor was sufficient to reduce binding of Gac by 2.6-fold, while a 25-fold excess of that competitor completely diminished the shift. Inclusion of 10-fold excesses of the empty pCR2.1 amplicon, *flaB*, or *ospAB* unlabeled DNAs exhibited lower levels of competition for Gac binding with none resulting in a fold change of greater than 1.3-times (Fig 3B, lanes 5, 6, and 7, respectively). However, 25-fold excess of each of those unlabeled DNAs competed Gac away from the labeled B31 *ospC* labeled probe (Supplemental Figure S1). These results indicate that Gac binds to the B31 *ospC* operator with a greater affinity than to the *flaB, ospAB* or pCR2.1 competitor sequences, but has a lower affinity for the B31 *ospC* operator than for the Ip89 *ospC* operator.

Using labeled *ospC* operator from strain B31 as the EMSA probe, truncated segments of that DNA region were used as competitors to narrow down the sequences that bound Gac (Fig 4). As above, 10- and 25-fold excesses of unlabeled probe DNA competed Gac away from the B31 *ospC* operator (Fig 4, lanes 3 and 4). No fragment competed as well as the unlabeled probe, although some competed more efficiently than others (Fig. 4, lanes 5-11). Notably, competitor D was more effective that the similarly-sized competitor C, indicating that Gac-binding preferentially involves nucleotides present in competitor D but absent from competitor C. The smaller unlabeled DNAs did not compete as well as the longer DNAs, suggesting that Gac interacts with extended and/or separate sequences in the B31 *ospC* 5’ region. Binding of Gac to DNA has been demonstrated to involve direct interactions with 40-50bp and additional associations with neighboring nucleotides (52, 55-57).

**Figure 4.**
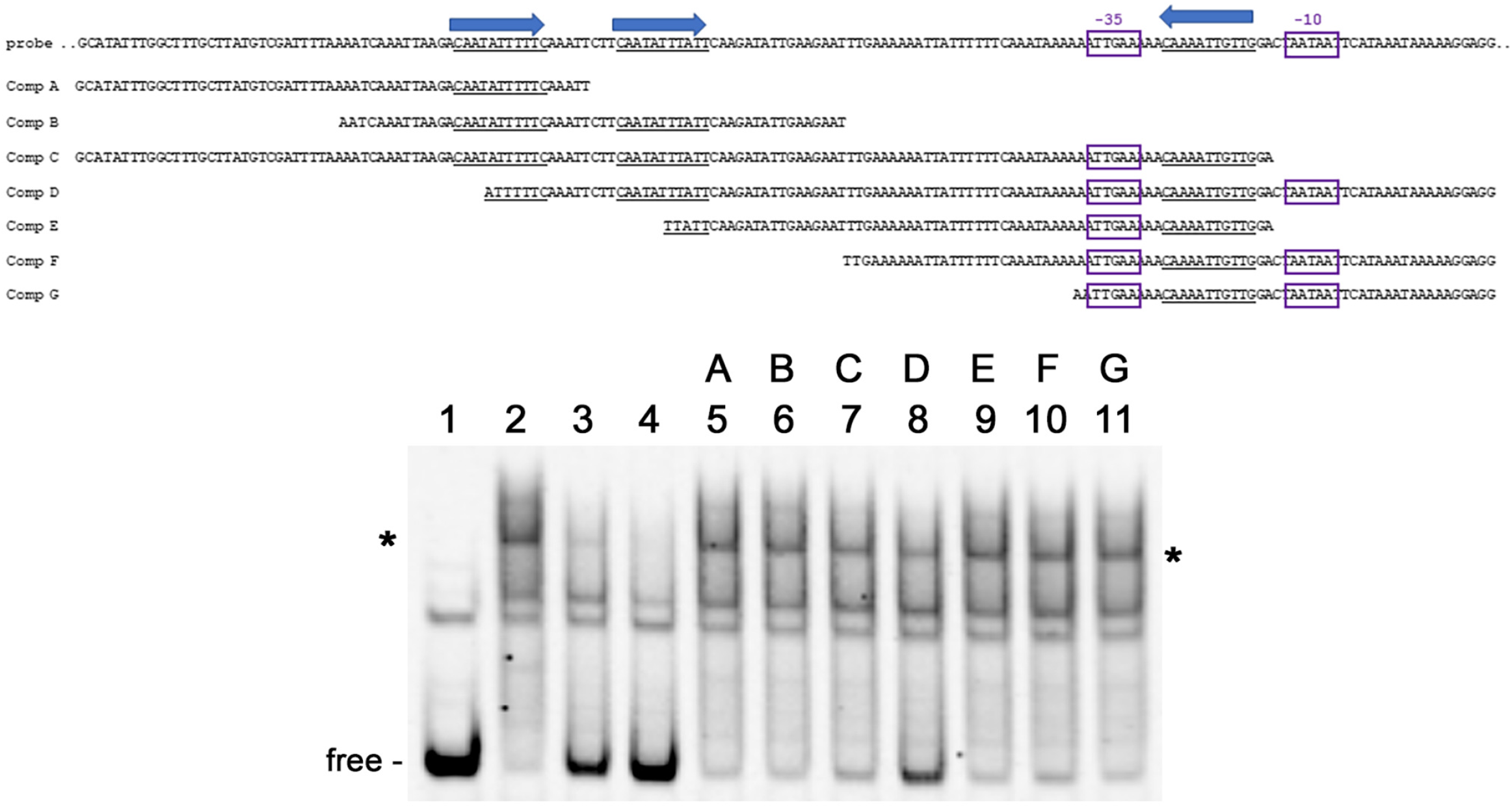
Mapping Gac binding sites in B31 *ospC* 5’ DNA by EMSA with purified recombinant Gac and unlabeled competitor DNA fragments. Sequences of the probe and competitors are shown above the EMSA, with the 11bp repeats indicated by blue arrows. Shifted DNAs are indicated with asterisks. Unoccupied DNAs are labeled “free”. Labeled B31 operator (10 nM) without (lane 1) and with 25 nM Gac (lanes 2-11). Lane 2: no added competitor. Lane 3: plus 10x excess unlabeled B31 *ospC* operator probe. Lane 4: plus 25x excess unlabeled B31 *ospC* operator probe. Lanes 5-11: plus 10x unlabeled competitors A through G, respectively.

### Altered *ospC* expression in *gac* mutant *B. burgdorferi*

To assess the effects of Gac in live *B. burgdorferi, gac* mutant strain CKO-1, strain CKO-1(pBLS820) (CKO-1 complemented with wild-type *gac*), and strain NGR (an isogenic wild-type *gac* clone) were cultured to mid-exponential phase at 23 °C, then diluted into fresh media and incubated to mid-exponential phase at 35 °C (6, 32). Immunoblot analyses of lysates of the 23 and 35°C cultures using OspC-specific antibodies found that all three strains exhibited differential expression of OspC protein, with greater amounts of OspC being synthesized at 35 °C (Fig. 5). However, Gac-deficient strain CKO-1 produced 6.4 to 23-times more OspC than did NGR at 23°C, consistent with Gac being a repressor of OspC synthesis. Levels of OspC in those two strains were very similar at 35°C, ranging between 0.92 to 1.4 times more OspC in CKO-1 than in NGR. Complementation of CKO-1 with a wild-type *gac* gene on pBLS820 reduced OspC levels by 1.6 to 6.9-times at 23°C, and by 1.2 to 4.3-times at 35°C. These results suggest that relief of Gac repression is a major contributor to the induction of OspC.

**Figure 5.**
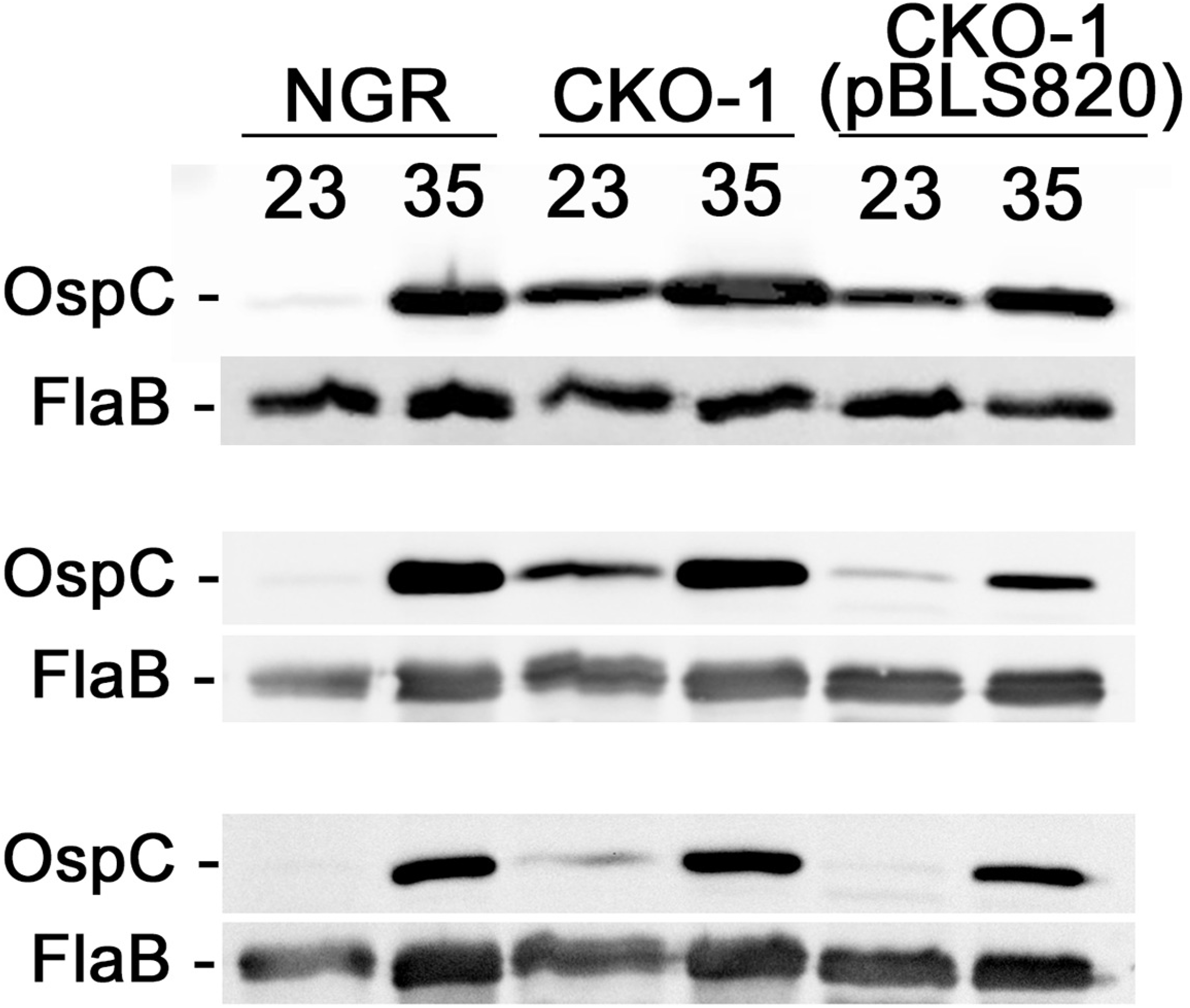
Triplicate western blot analyses of strains NGR, CKO-1, and CKO-1(pBLS820) that were cultured to mid-exponential phase at 23 or 35°C, probed with monospecific antibodies against OspC and FlaB. Results of densitometric analyses are shown in Supplemental Table S2.

*B. burgdorferi* strains NGR and CKO-1 were next transformed with plasmids that carry transcriptional fusions between B31 *ospC* operator/promoter sequences and *gfp*. Plasmid pBLS761 carries the same operator/promoter sequence that was used as the EMSA probe in the above-described studies, while pBLS764 lacks the DNA 5’ of the *ospC* promoter. Each transformant was cultured at 35°C to mid-exponential phase, then bacterial GFP content was assessed by flow cytometry (Fig. 6). CKO-1(pBLS761) produced substantially greater levels of GFP than did NGR(pBLS761), consistent with the observed levels of OspC protein and transcript described above.

**Figure 6.**
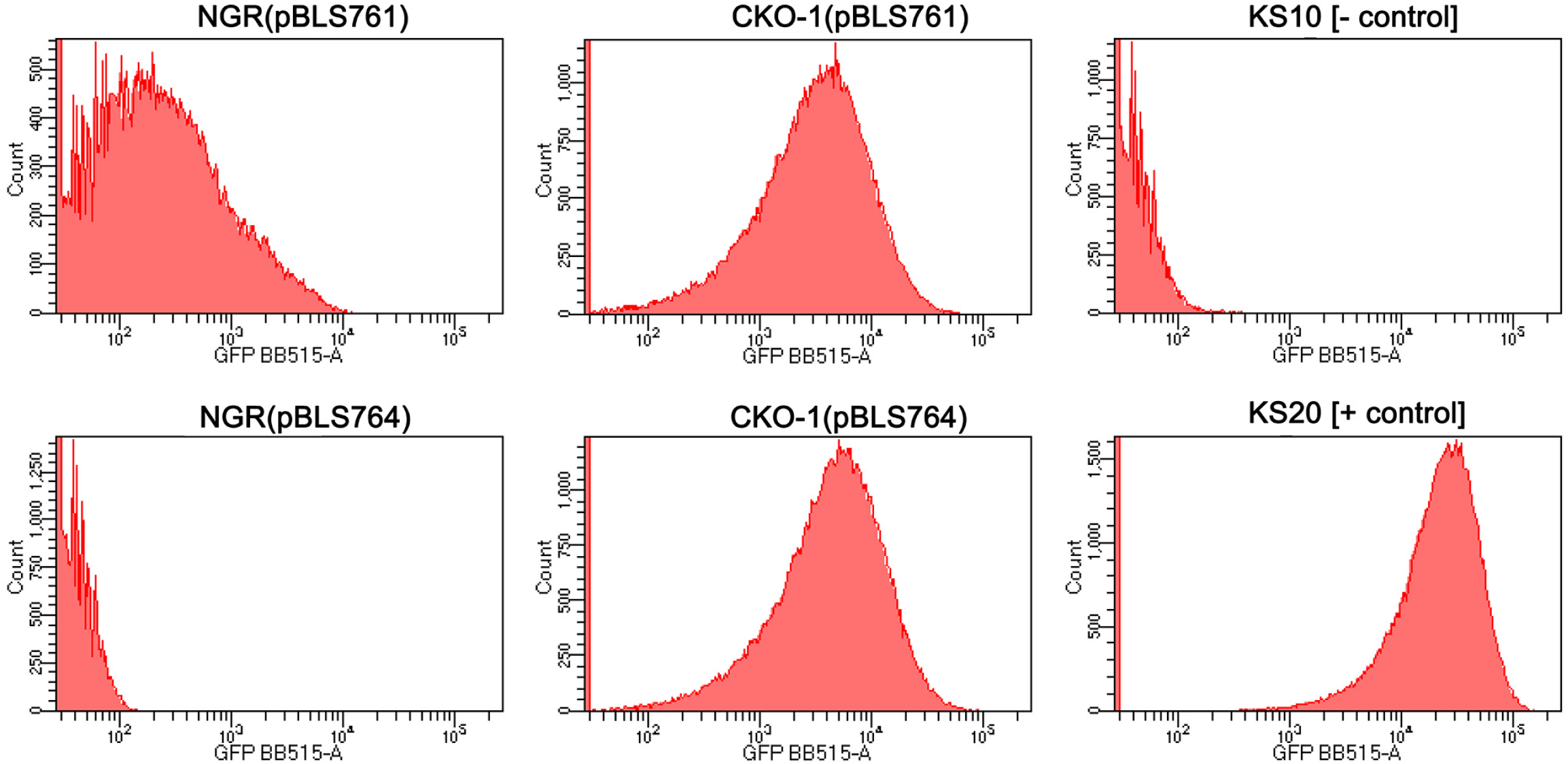
Flow cytometric analyses of green fluorescent protein (GFP) levels. Each panel represent approximately 100,000 bacteria that were cultured to mid-exponential phase (approximately 5×10^7^ bacteria/ml) at 35°C. Isogenic NGR and *gac* mutant CKO-1 contained transcriptional fusions of the *ospC* promoter, with an without the operator (pBLS761 and pBLS764, respectively), fused to *gfp*. Negative control strain KS10 carries promoterless *gfp* on pBLS590, and positive control strain KS20 carries the constitutive *erpAB* promoter fused to *gfp* on pBLS599 (73).

NGR(pBLS764), which lacks the *ospC* operator, did not produce GFP (Fig. 6). In contrast, CKO-1(pBLS764) produced GFP at levels comparable to CKO-1(pBLS761). We conclude that absence of Gac relieves transcriptional repression of the *ospC* promoter, and that the *ospC* operator is also required for efficient transcription from the *ospC* promoter if Gac is present. Those results suggest that one or more additional factors interact with *ospC* operator DNA to inhibit Gac binding to the *ospC* promoter. Studies are ongoing to examine that hypothesis.

### *B. burgdorferi* regulates transcription of *gac*

Since *ospC* is induced in *B. burgdorferi* cultured at 35°C, we assessed *ospC* and *gyrA/gac* transcript levels over time in cultures of wild-type *B. burgdorferi* (Fig. 7). Levels of *ospC* transcript increased as the cultures grew through exponential into stationary phase. The two *gyrA/gac* primer sets yielded different Q-RT-PCR results for each pair. Since the 5’ primers detected mRNA that only encoded *gyrA*, whereas the 3’ primers detected both *gyrA* and *gac* mRNAs, the relative concentration of *gac* mRNA was approximated by subtracting the results of 5’ Q-RT-PCR from results with the 3’ primer set. The concentration of Gac-encoding transcript was greatest during lag and early exponential phases, then declined as the cultures matured and entered stationary phase. Those changes were opposite of *ospC*, and are consistent with the above-described results that indicate Gac is a repressor of *ospC*.

**Figure 7.**
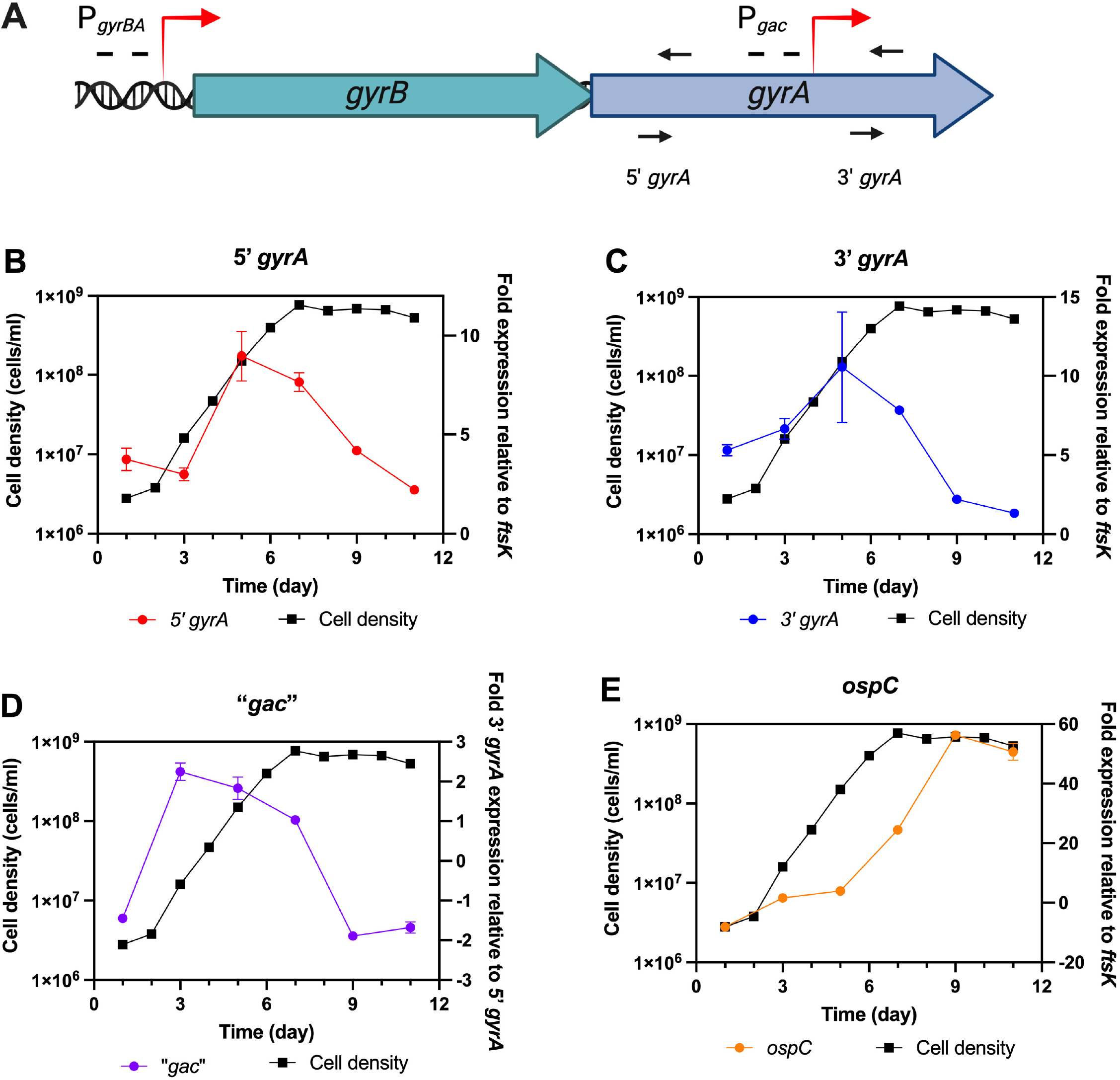
Growth curves and quantitative-RT-PCR analyses of selected transcripts of *B. burgdorferi* B31, cultured at 35°C. Bacterial cell densities are indicated by black squares and lines. Relative transcript levels are indicated by red circles and lines. **(A)** Schematic of the locations of amplicons used to determine relative expression of the 5’ and 3’ portions of *gyrA*. The 5’ primer pair amplified only transcripts arising from the promoter 5’ of *gyrB*, while the 3’ primer pair amplified transcripts arising from both the promoter 5’ of *gyrB* and within the *gyrA* ORF. **(B)** Results from the 5’ segment of *gyrA*. **(C)** Results from the 3’ segment of *gyrA*. **(D)** Results from the *gyrA* 3’ segment minus the *gyrA* 5’. **(E)** Analyses of *ospC* transcript levels.

### Gac exhibits affinity for the telomere of *B. burgdorferi* B31 plasmid lp17

*Borrelia* species contain both circular and linear DNAs, and there is considerable interest in the mechanisms by which the spirochetes maintain linear replicons (58-60). The Gac protein was initially identified through an affinity pull-down method similar to ours, using a bait sequence derived from a telomere of strain B31’s linear plasmid lp17 (53). Having found that Gac preferentially binds to *ospC* 5’ DNAs, we revisited the previous study. A labeled DNA probe with the same telomere-derived sequence was produced, then assessed by EMSA for ability to bind Gac in the presence of excess levels of various unlabeled DNAs. Recombinant Gac produced an EMSA shift with the labeled telomere probe, which was effectively competed away with 10-fold and 25-fold excesses of unlabeled telomere sequence (Fig. 8, lanes 2-4). Ten-fold excesses of unlabeled Ip89 or B31 *ospC* operator DNAs also competed Gac away from the labeled telomere DNA (Fig. 8, lanes 6 and 7, respectively). Unlabeled empty pCR2.1 amplicon partly competed for Gac binding, while neither the *flaB* nor the *ospAB* DNAs detectably competed (Fig. 8, lanes 5, 8, and 9, respectively). We conclude that Gac exhibits an affinity for the lp17 telomere region that is greater than for the control *ospAB* or *flaB* DNA sequences.

**Figure 8.**
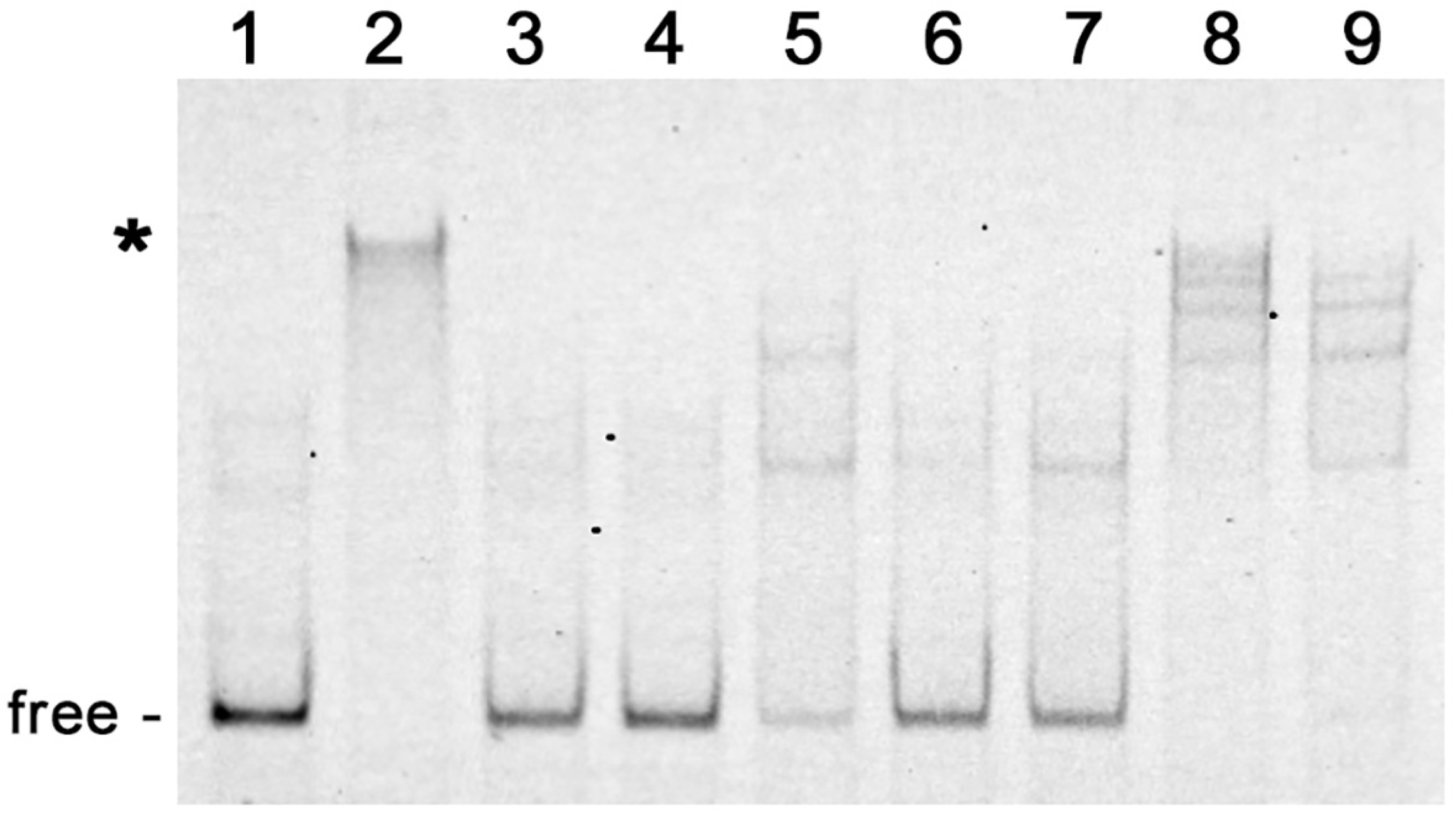
Representative EMSAs of purified recombinant Gac with labeled *ospC* lp17 telomere DNA and competitors. Shifted DNAs are indicated with asterisks. Unoccupied DNA is labeled “free”. All lanes contain 10 nM labeled lp17 telomere probe, without (lane 1) or with 25 nM Gac (lanes 2-9). Lane 2: no added competitor. Lane 3: plus 10x excess unlabeled lp17 telomere. Lane 4: plus 25x excess unlabeled lp17 telomere. Lane 5: plus 10x unlabeled pCR2.1 amplicon. Lane 6: plus 10x excess unlabeled Ip89 *ospC* operator. Lane 7: plus 10x excess unlabeled B31 *ospC* operator. Lane 8: plus 10x excess unlabeled *flaB* DNA. Lane 9: plus 10x unlabeled *ospAB* DNA.

## Discussion

These studies found that Gac exhibits preferential affinity for *ospC* promoter and operator DNA, and inhibits transcription of *ospC*. Studies in wild-type and Gac-deficient mutant bacteria found that absence of Gac resulted in substantially increased expression levels of both native OspC and *ospC* promoter :: *gfp* transcriptional fusions. Those results indicate that Gac is a major contributor to repression of OspC.

Deletion of the *ospC* operator did not alleviate Gac-dependent repression of transcription from the *ospC* promoter, suggesting that Gac binding to the promoter itself is sufficient to repress transcription. Additionally, the *ospC* operator is evidently necessary for relief of Gac-dependent transcriptional repression, mirroring results from previous studies that also found that the operator is essential for regulation of *ospC* transcription (42-44). Thus, it appears that one or more unknown factors interact with the *ospC* operator and somehow prevents repression by Gac (Fig. 9). On the hypothesis that Gac in wild-type *B. burgdorferi* cytoplasmic extracts might have outcompeted the predicted “factor X” for binding to the *ospC* operator/promoter probe used in the initial DNA affinity pull-downs, we recently performed similar analyses using extract from Gac-deficient mutant CKO-1. Salt fractionation and SDS-PAGE revealed two proteins with sizes different from Gac (our unpublished results). We are currently examining whether these might be the hypothesized “factor X” anti-repressor.

**Figure 9.**
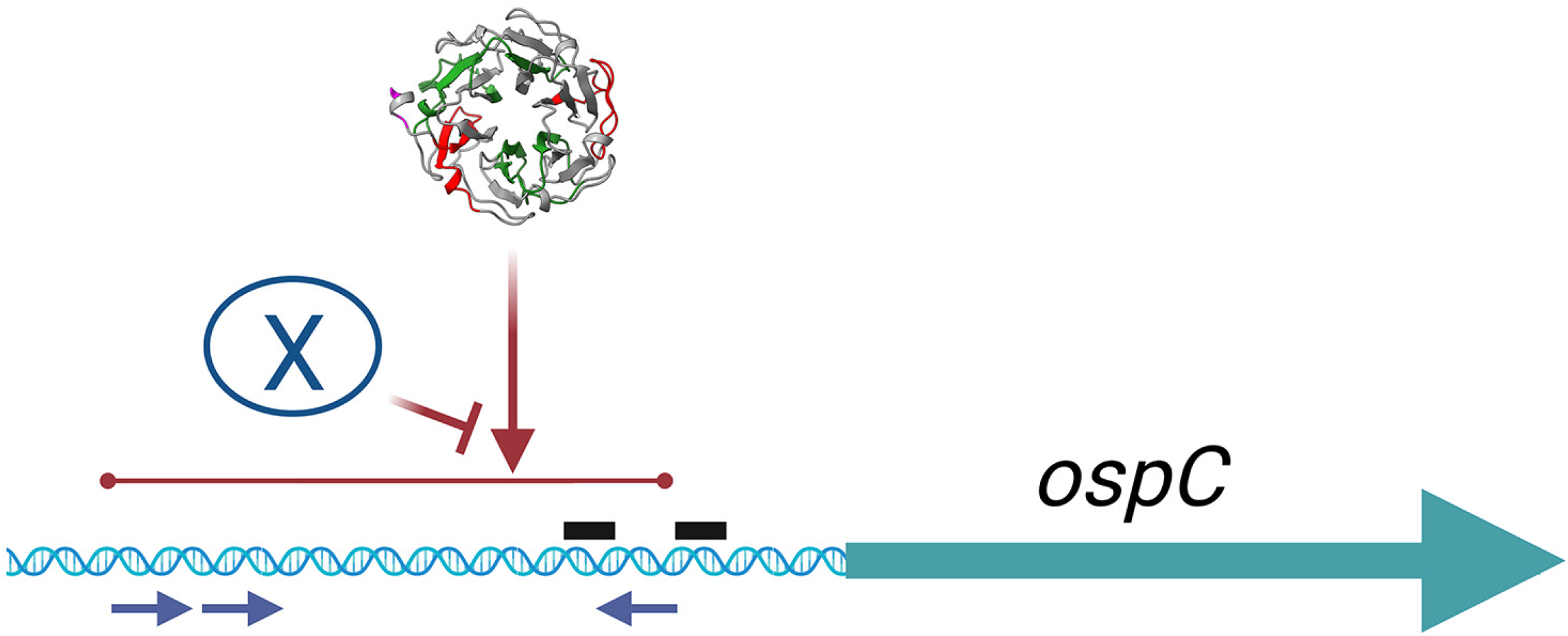
Graphic summary of these results. Gac binds at or near the 11bp repeated sequences in the *ospC* promoter and operator, and inhibits transcription from the *ospC* promoter. Deletion of the *ospC* operator resulted in ablation of transcription from the *ospC* promoter, as has also been noted previously (42-44). Those data indicate that another factor, “X”, interacts with the *ospC* operator and is required to activate transcription. In the absence of Gac, the *ospC* promoter is transcribed at high levels, indicating that “X” competes against Gac repression. X may be one or more protein or other factor. The 11bp repeat sequences are indicated by blue arrows beneath the DNA. The *ospC* promoter -10 and -35 sequences are indicated by black bars above the DNA.

An earlier study found that inhibition of *B. burgdorferi* DNA gyrase function by addition of coumermycin resulted in increased production of OspC protein (44). Considering the effect of Gac on *ospC* expression, it will be important to determine whether the effect of coumermycin is due to changes in DNA topology at the *ospC* operon, changes in levels of Gac or other regulatory proteins, or a combination of those and, perhaps, other factors.

Our studies found that Gac has elevated affinity for three sites in the *B. burgdorferi* genome – the *ospC* operator, the *ospC* promoter, and the telomere of lp17 – yet can also possesses lesser affinity for other DNA sequences. Gac wraps DNA around itself, forming 180-220° bends (52). Gac complements some features of the *E. coli* nucleoid-associate protein HU when expressed in that species (53, 54). Taken together, these data suggest that Gac may be a nucleoid-associated protein (“histone-like protein”). It is not uncommon for nucleoid-associated proteins to also influence transcription (61-63). Indeed, the antirepressor of *B. burgdorferi erp* operon transcription, EbfC, meets the criteria of being an NAP (64).

The ability of Gac to preferentially bind to certain DNA sequences raises the possibility that GyrA, alone or as part of DNA gyrase, might also have a preference for these sequences. We are currently investigating this possibility, as it could affect gene expression and aspects of DNA supercoiling.

At this time, it is not known whether production of a separate Gac protein from the 3’ end of the *gyrA* ORF is unique to *B. burgdorferi*. The possibility that *gyrA* can encode two distinct proteins has never been addressed for the vast majority of other bacterial species. It would be worth examining other *gyrA* genes for presence of potential ribosome binding sites and start codons near the beginning of the C-terminal domain.

In conclusion, our investigations found that *B. burgdorferi* Gac has elevated affinity for *ospC* operator/promoter DNAs. Gac repressed *ospC* transcription in vitro and in live spirochetes. Levels of *ospC* and *gac* transcripts changed in opposite directions during cultivation, consistent with *ospC* repression by Gac. Analyses of mutants indicate that the *ospC* operator is required to alleviate Gac repression of the *ospC* promoter, implying that one or more additional factors interact with the *ospC* operator to prevent repression by Gac. The Gac protein also bound with elevated affinity to an lp17 telomere, suggesting that it might play a role in linear DNA maintenance. Our observations, and those of others (52, 53), indicate that Gac functions like a nucleoid associated protein that exerts effects on gene regulation and genomic structure.

## Materials and Methods

### Bacteria, plasmids, and culture conditions

*B. burgdorferi* sensu stricto strain B31 is the type strain, and was isolated from an *Ixodes scapularis* tick that was collected on Shelter Island, New York (65, 66). Infectious clone B31-MI-16 was used to prepare cytoplasmic extracts for identification of *ospC* operator-binding proteins (67). *B. garinii* strain Ip89 was isolated from an *I. persulcatus* tick that was collected in central Eurasia (68, 69).

*Borrelia* spp. strains were cultured in BSK-II broth at either 23 or 35°C (70). Temperature shift cultivation was performed by diluting a 23°C culture 1:100 into fresh medium, then incubating at 35°C (32). Culture densities were determined with a Petroff-Hausser counting chamber and dark field microscopy.

Analyses of the effects of Gac on OspC expression used a ∆*gac* clone of strain B31, CKO-1, and an isogenic control strain, NGR (54). The overlap of the *gac* ORF with the essential *gyrA* ORF precludes outright deletion of the *gac* locus. For that reason, Knight et al. (54) produced *gac* defective strain CKO-1 by an allelic exchange mutation that changed the two potential *gac* start codons from ATG to CTT and ATT (leucine and isoleucine, respectively) and the ribosome binding site from AGGAGG to AGAAGA. The mutant does not produce a Gac protein, although it does produce normal levels of GyrA (54). The *gac* mutations were selected by simultaneous introduction of point mutations into the adjacent *gyrB* gene, which confer resistance to coumermycin (54, 71). The CKO-1 *gac* mutant exhibits growth patterns that are physiologically indistinguishable from the wild-type parent or from NGR, an isogenic coumermycin-resistant *gyrB* strain (54).

Plasmid pBLS820 was custom produced by GenScript (Piscataway, NJ) by cloning the *B. burgdorferi gac* open reading frame, ribosome-binding site, and promoter into vector pBSV2-G (Supplemental Fig. S2) (72). *B. burgdorferi* CKO-1 was transformed by electroporation, and transformants were selected by culture in 150 µg/ml gentamicin.

Plasmid pBLS761 contains the B31 *ospC* promoter and operator in a transcriptional fusion with *gfp*. Plasmid pBLS764 is identical to pBLS761 except that it lacks the *ospC* operator. Both are based on pBLS740, which was produced for this work by cloning the *gfp* gene of pBLS590 between the BamHI and HindIII sites of vector pKFSS1 (73, 74). The *ospC*-derived DNAs used to make pBLS761 and pBLS764 were inserted between the KpnI and BamHI sites of pBLS740, custom synthesized by GenScript. *B. burgdorferi* strains NGR and CKO-1 were individually transformed with plasmids pBLS761 and pBLS764, to produce strains NGR(pBLS761), NGR(pBLS764, CKO-1(pBLS761), and CKO-1(pBLS764). Two of our previously-described *B. burgdorferi* strains were used as GFP expression controls: KS10, which carries a promoterless *gfp* on pBLS590, and KS20, which carries the constitutively-expressed minimal *erpAB* promoter driving *gfp* on pBLS599 (73). Plasmids pBLS590 and pBLS599 confer resistance to kanamycin. *B. burgdorferi* containing pBLS590 or pBLS599 were cultured in media that contained 200 µg/ml kanamycin, while spirochetes that contain pBLS761 or pBLS764 were cultured in media with 50 µg/ml streptomycin (73-76).

### Cytoplasmic extracts for EMSA

*B. burgdorferi* ss B31-MI-16 was cultured at 35°C to mid-exponential phase (approximately 5×10^7^ bacteria/ml). Bacteria were harvested by centrifugation, washed with phosphate buffered saline (PBS), and resuspended in a minimal volume of 50mM Tris (pH=7.6), 10mM EDTA, 10% (w/vol) sucrose. The suspension was frozen in a dry ice/ ethanol bath, then allowed to thaw in an ice-water bath. NaCl, dithiothreitol, and lysozyme were added to final concentrations of 140 mM, 1 mM, and 0.4 mg/ml, respectively, and incubated on ice for 5 min. The cell mixture was subjected to four additional freeze-thaw cycles. Cellular debris was pelleted by centrifugation, and the supernatant was aliquoted and stored at -80°C (77).

### Electrophoretic mobility shift assays (EMSAs)

Labeled and unlabeled nucleic acids are described in Supplementary Table 1. Fluorescently tagged IRDye800 labeled probes were utilized for EMSAs. For each EMSA binding reaction, a labeled probe was incubated for 15 min at room temperature with protein and unlabeled competitor DNA as necessary, in EMSA buffer (50mM Tris-HCl, 25mMKCl, 10% glycerol (vol/vol), 0.01% Tween 20, 100nM dithiothreitol, and 1mM phenylmethanesulfonyl fluoride (PMSF). Protein-nucleic acid mixtures were then loaded into either 6% or 10% Novex TBE gels (Thermo Fisher). Gels were pre-run for at least 30 min at 100V in 0.5x TBE buffer before sample loading. Following the addition of the samples, electrophoresis was conducted for up to 1.5 h at 100V.

Initially, a fragment of *B. garinii* Ip89 DNA, from 5’ of the *ospC* operator through the *ospC* promoter, was amplified from genomic DNA by PCR with oligonucleotide primers 1021 and 1022 (Supplementary Table 1). The resulting amplicon was cloned into pCR2.1 (Invitrogen) to create pBLS756. Fluorescently-tagged IP89 *ospC* operator was produced by PCR of pBLS756 using IRDye800-tagged M13-Reverse and unlabeled M13-Forward primers (Supplementary Table 1). The probe was incubated with cleared borrelial cytoplasmic extract in EMSA buffer for 15 min at room temperature with 10ng poly-dI-dC, then subjected to electrophoresis. These exploratory EMSAs were performed twice.

Other EMSAs used purified recombinant Gac protein, labeled *ospC* operator/promoter probes, and unlabeled DNA competitors. Unless noted otherwise, all EMSAs were performed a minimum of three times.

In addition, a sub-telomeric sequence of strain B31 lp17 was amplified from total genomic DNA by PCR with oligonucleotides TL16h and TL16g as previously described (53). The resulting PCR product was cloned into pCR2.1 (Invitrogen) and maintained in *E. coli* TOP-10. The amplicon was completely sequenced on both strands to ensure fidelity. Fluorescently tagged telomere probe was produced by PCR using IRDye800 tagged M13-Reverse and unlabeled M13-Forward primers.

Densitometric analyses of EMSAs was performed using Image Lab 6.1 (Bio-Rad) software. Lanes and bands were added manually and subsequently analyzed to generate the lane percentage totals for each band in a lane. Protein-DNA shifts were considered as the entire region above the free DNA in each lane. The free DNA was determined for each lane by analyzing the band percentage values. For each EMSA, the free DNA percentage values for all lanes were normalized to the probe-only control band percentage value. This set the free DNA percentage of the probe-only control as equal 100%. The normalized free DNA values were then subtracted from 100% to calculate the percent shifted DNA in each lane. Fold changes in DNA binding of EMSA competition assays were calculated by dividing the percent shifted in competition lanes by the percent shifted of the protein-probe only lane.

### DNA affinity pull-down

Proteins that bind *ospC* operator DNA were purified from B31-MI-16 cytoplasmic extracts as previously described (78). Briefly, bacteria of a mid-exponential phase (approximately 5×10^7^ bacteria/ml) culture were harvested, washed, and frozen at -80°C. Bacterial pellets were then thawed on ice, and subjected to three additional rounds of freezing at -80°C / thawing on ice. Bacterial lysates were resuspended in a minimal volume of BS/THES buffer (44.3% THES buffer, 20% BS buffer, 35.7% water). THES Buffer consists of 50mM Tri HCl (pH 7.5), 10mM EDTA, 20% sucrose (w/vol), 140mM NaCl, 0.7% (vol/vol) Protease Inhibitor Cocktail II (Sigma), and 0.1% (vol/vol) Phosphatase Inhibitor Cocktail II (Sigma). BS buffer consists of 50mM HEPES (pH=7.5), 25mM CaCl_2_, 250mM KCl, and 60% (vol/vol) glycerol. Suspended bacteria were lysed by sonication, on ice. Cellular debris was pelleted by centrifugation, and the supernatant was divided into aliquots.

Recombinant plasmid pBLS756, which contains the *B. garinii* Ip89 *ospC* operator/promoter sequence, was used as template to produce a biotin-tagged amplicon by PCR with biotin-tagged M13-Reverse and unlabeled M13-Forward primers (Supplementary Table 1). The probe was incubated with cleared borrelial cytoplasmic extract in EMSA buffer for 15 min at room temperature with 10ng poly-dI-dC, then subjected to electrophoresis. The bait DNA was bound to streptavidin Dynal beads (Invitrogen), then washed with BS/THES buffer. Cytoplasmic extracts were incubated with the DNA-bound beads, and washed extensively with BS/THES. Bound proteins were eluted by washing with buffer containing increasing NaCl concentrations to 0.2, 0.3, 0.5, 0.75, and 1.0 M. Fractions were analyzed by SDS-PAGE, and proteins were stained with SYPRO-Ruby (Molecular Probes, Eugene, OR). A protein band that eluted at 0.3 and 0.5 M NaCl was cut out. The samples were treated with trypsin and assayed by LC-MS-MS at the University of Kentucky Proteomics Core. Results were compared with the *B. burgdorferi* strain B31-MI sequence (59, 60) using Mascot (Matrix Science, Boston).

### Recombinant Gac protein

The B31 *gac* ORF was cloned into pET101, then sequenced to confirm fidelity. C-terminally His-tagged Gac was expressed in *E. coli* Rosetta II upon induction with 1 mM of isopropyl-β-D-thiogalactopyranoside (IPTG) for 1 hour at 37°C. Bacteria were collected by centrifugation, and the pellet was frozen at -80°C. The bacterial pellet was then thawed on ice, resuspended in phosphate buffered saline (PBS) containing 10 mM imidazole, and bacteria lysed by sonication. The sonicate was subjected to centrifugation, then recombinant protein purified from the soluble fraction using HisPur Ni-NTA columns (Thermo Scientific). Purified proteins were dialyzed against EMSA buffer. Protein concentrations were determined by Bradford assay and purity assessed by SDS-PAGE with Coomassie brilliant blue staining.

### Immunoblot analyses

Cultures of strains NGR, CKO-1, and CKO-1(pBLS820) were harvested at mid-exponential phase by centrifugation, then split into aliquots for immunoblot assays and RNA extraction. Pellets for immunoblot assays were frozen at - 80°C, subsequently thawed at room temperature, then resuspended in 10% SDS, boiled for 5-minutes, and vigorously vortexed. The resulting lysates were stored at 4°C. Prior to electrophoresis, samples were preheated to 42° before adding to 2x sodium dodecyl sulfate (SDS)-page loading dye (63mM Tris pH7.5, 18.75% [mass/vol] glycerol, 10% [vol/vol] 2-mercaptoethanol, 4% [mass/vol] SDS, and .5% [mass/vol] bromophenol blue). Separation by SDS-polyacrylamide gel electrophoresis was followed by electrotransfer to nitrocellulose membranes and blocking with 5% nonfat dried milk in Tris-buffered saline-Tween-20 (20 mM Tris [pH 7.5], 150 mM NaCl, 0.05% [vol/vol] Tween-20). The membranes were then incubated with murine monoclonal antibodies directed against FlaB or OspC (79, 80). After incubation with primary antibodies, membranes were incubated with goat anti-mouse IgG conjugated to IRDye800 (LI-Cor). Immunoblots were imaged with a ChemiDoc MP (Bio-Rad, CA) and densities analyzed with background compensation using Image-Lab software (Bio-Rad). All immunoblots were performed a minimum of three times.

### Flow cytometric analysis of GFP expression

Transcriptional activity of *ospC* promoter::*gfp* reporter constructs was assayed essentially as we previously described (73). Briefly, cultures of s NGR(pBLS761), NGR(pBLS764, CKO-1(pBLS761), CKO-1(pBLS764), KS10, and KS20 were grown to mid-exponential phase (approximately 5×10^7^ bacteria per ml) in BSK-II broth at 35°C. Bacteria were collected by centrifugation, washed twice with phosphate-buffered saline (PBS), then resuspended in PBS at concentrations of approximately 10^6^ bacteria per ml. Aliquots of each bacterial suspension were analyzed using a FACSymphony flow cytometer (Becton Dickinson, San Jose, Calif.), with excitation and detection wavelengths of 488 and 530 nm, respectively. Approximately 100,000 events were measured for each flow cytometry analysis.

### Quantitative reverse transcription PCR (Q-RT-PCR)

*B. burgdorferi* was cultured at 35ºC, and 2 ml aliquots of bacteria were taken 1-, 3-, 5-, 7-, 9-, and 11-days post-inoculation. Bacteria were pelleted, then washed two times with PBS and frozen at - 80ºC. Frozen pellets were later resuspended in 60ºC TRIzol (Thermo Fisher), and RNA was extracted using Qiagen Mini RNA kits. Residual genomic DNA was depleted using on-column Turbo DNase (Thermo Fisher). The quality (RIN > 7) and quantity of the RNA were determined using an Agilent 2100 Bioanalyzer and Agilent 6000 Nano chips. Purified RNA stocks were stored at -80ºC.

Complementary DNA (cDNA) for qPCR was produced using the iScript gDNA Clear cDNA synthesis kits (Bio-Rad). The cDNA stocks were diluted 1:100 into nuclease-free water for use as template. Quantitative PCR was carried out using iTaq Universal SYBR Green Supermix (Bio-Rad) and a Bio-Rad CFX96 Touch Real-time PCR thermocycler. For all reactions, cycles involved melting at 95ºC for 10s, annealing at 55ºC for 10 s, and extension at 72ºC for 30s. Each run was performed with technical triplicates of each reaction as well as no-template control reactions that lacked template. Melting curves were performed with each run to validate production of specific amplicons. Every reaction was tested with a reverse transcriptase-negative template to validate the absence of contaminating gDNA. Cq values were normalized to *ftsK*, a gene whose expression is stable across different growth phases (81), using the ΔCq method, and fold-difference was determined by the function 2^− Δ Cq^ (82).

## Acknowledgements

This work was funded by NIH grant 3R01AI144126-03S1. We thank Jessamyn Moore for technical assistance and constructive comments; Scott Samuels for providing *B. burgdorferi* strains NGR and CKO-1, and for helpful comments on these studies; Haining Zhu and Jing Chen for assistance with mass spectrometric analyses; and Jennifer Strange, Greg Bowman, and Cuiping Zhang for flow cytometry analyses. Figure 9 was produced using BioRender.com.

## Supplemental Tables and Figures

**Supplemental Table S1.**
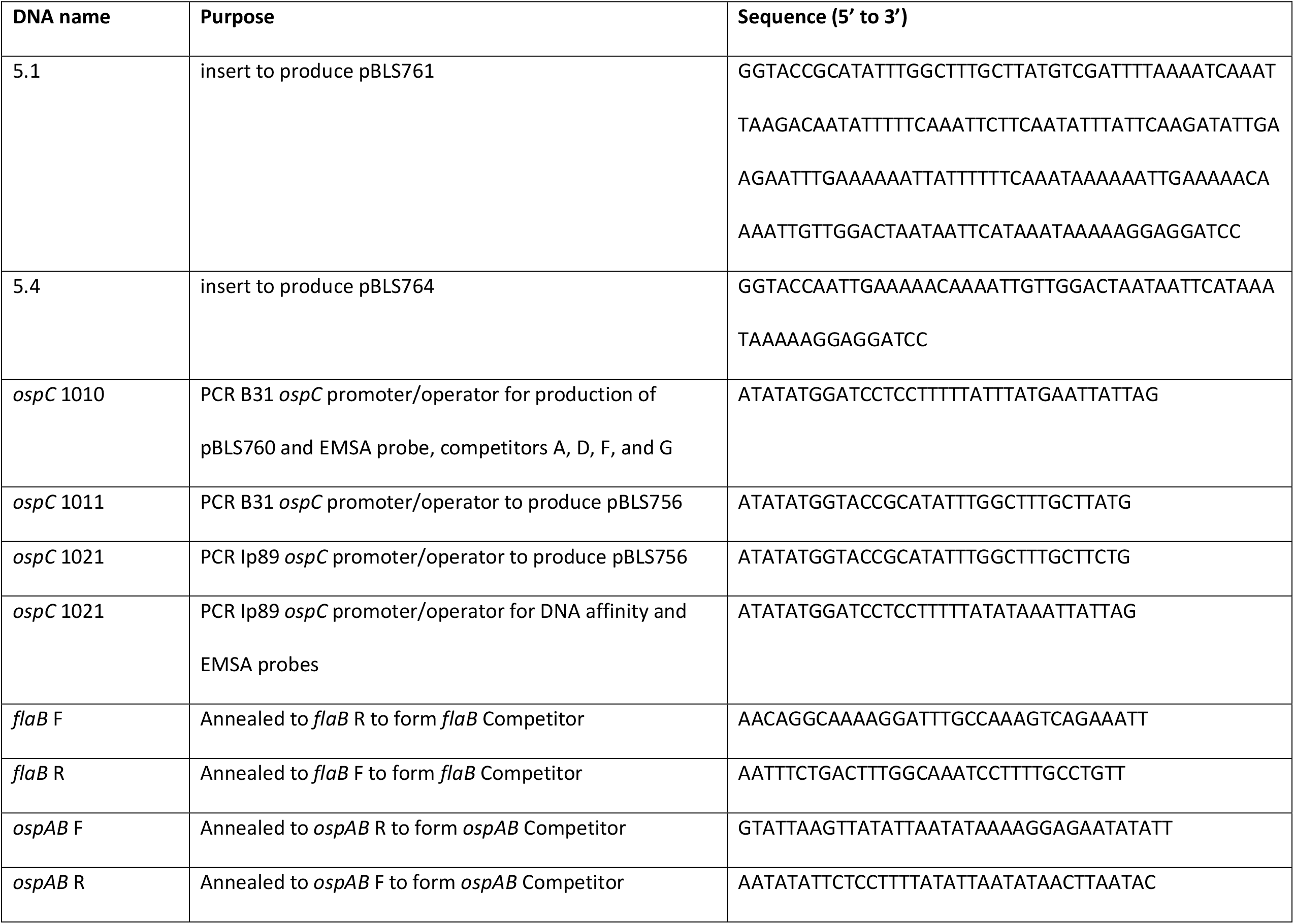
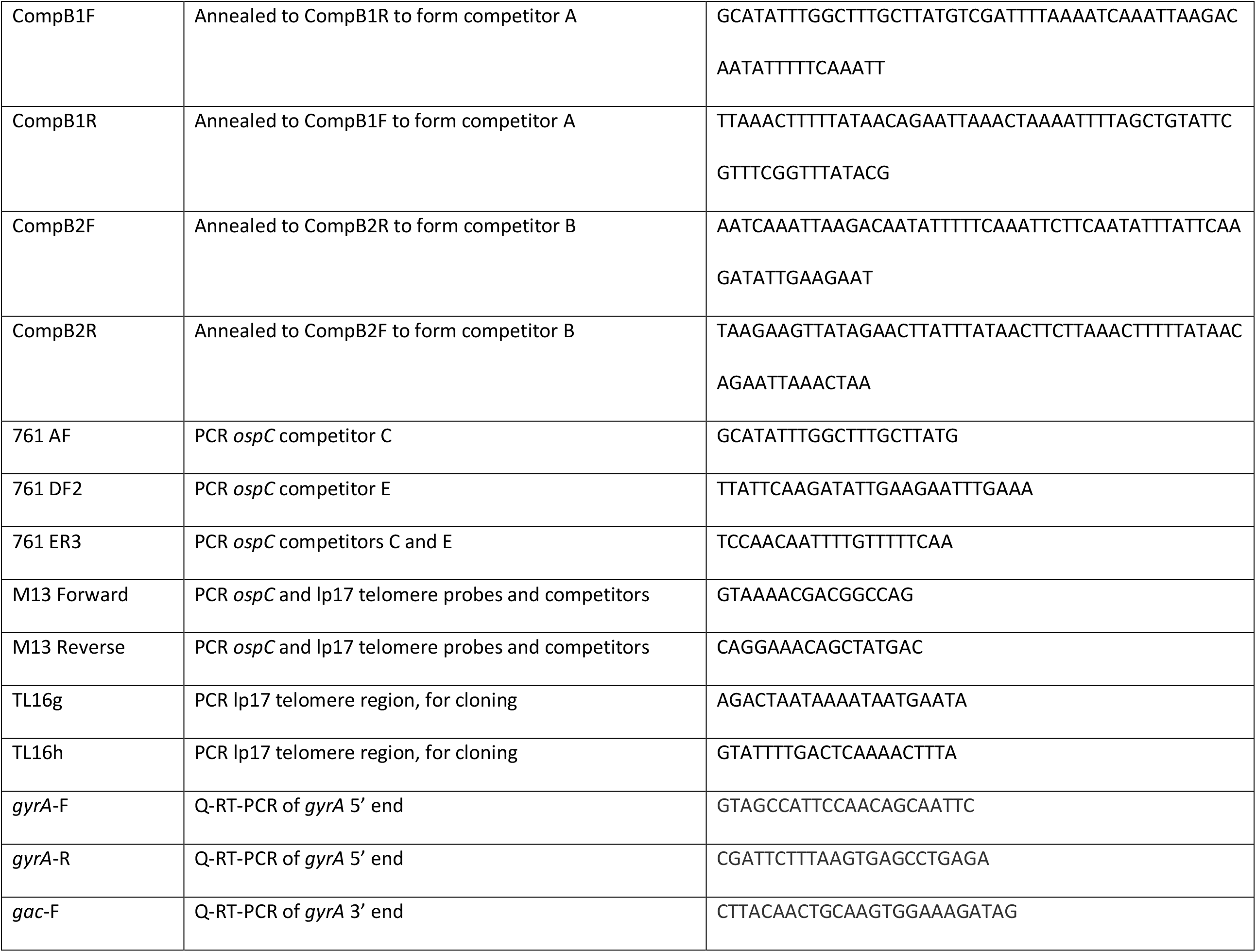
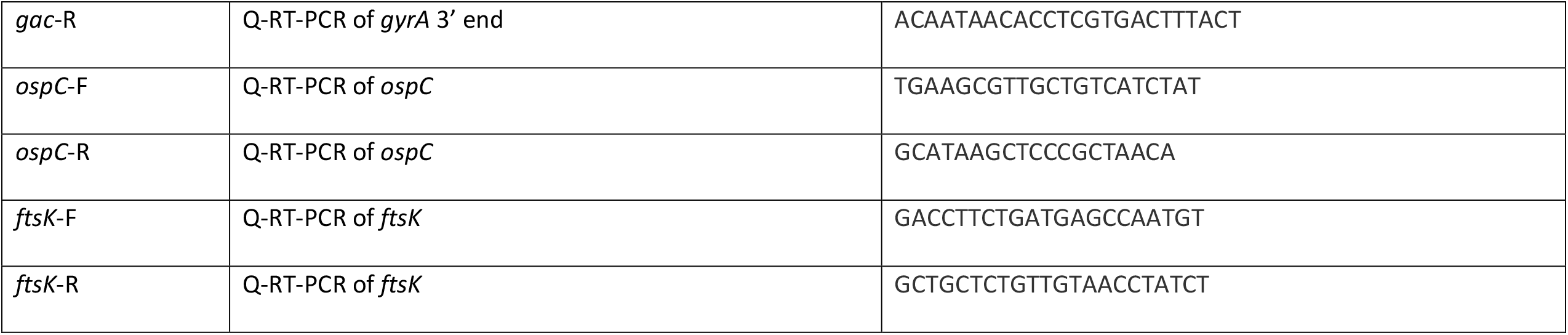
Sequences of synthetic DNAs and oligonucleotide primers used in this study

**Supplemental Table S2.** Relative changes in OspC protein levels in analyses of *B. burgdorferi* NGR, CKO-1, and CKO-1 (pBLS820), as shown in Figure 5. The immunoblots were imaged with a ChemiDoc MP (Bio-Rad, CA), and densities analyzed with background compensation using Image-Lab software (Bio-Rad). **A**. relative differences between strains at the same temperature. For example, in western 1, CKO-1 produced 23x more OspC than did NGR at 23°C. **B**. relative changes of the same strain at 23oC vs. 35oC. For example, in western 1, NGR produced 32x more OspC after shift to 34°C.

**A.**
| Western | Strain | Temperature |  |
| --- | --- | --- | --- |
|  |  | 23°C | 35°C |
| 1 | CKO-1 vs NGR | 23 | 1.4 |
|  | CKO-1 (pBLS820) vs NGR | 14 | 1.1 |
|  | CKO-1 vs CKO-1 (pBLS820) | 1.6 | 1.2 |
| 2 | CKO-1 vs NGR | 17 | 0.92 |
|  | CKO-1 (pBLS820) vs NGR | 2.5 | 0.22 |
|  | CKO-1 vs CKO-1 (pBLS820) | 6.9 | 4.3 |
| 3 | CKO-1 vs NGR | 6.4 | 1.1 |
|  | CKO-1 (pBLS820) vs NGR | 1.5 | 0.69 |
|  | CKO-1 vs CKO-1 (pBLS820) | 4.2 | 1.2 |

**B.**
| Western | Strain | Change when Shifted from<br>23°C to 35°C |
| --- | --- | --- |
| 1 | NGR | 32 |
|  | CKO-1 | 2.0 |
|  | CKO-1 (pBLS820) | 2.6 |
| 2 | NGR | 72 |
|  | CKO-1 | 3.9 |
|  | CKO-1 (pBLS820) | 6.2 |
| 3 | NGR | 57 |
|  | CKO-1 | 10 |
|  | CKO-1 (pBLS820) | 26 |

**Supplemental Figure S1.**
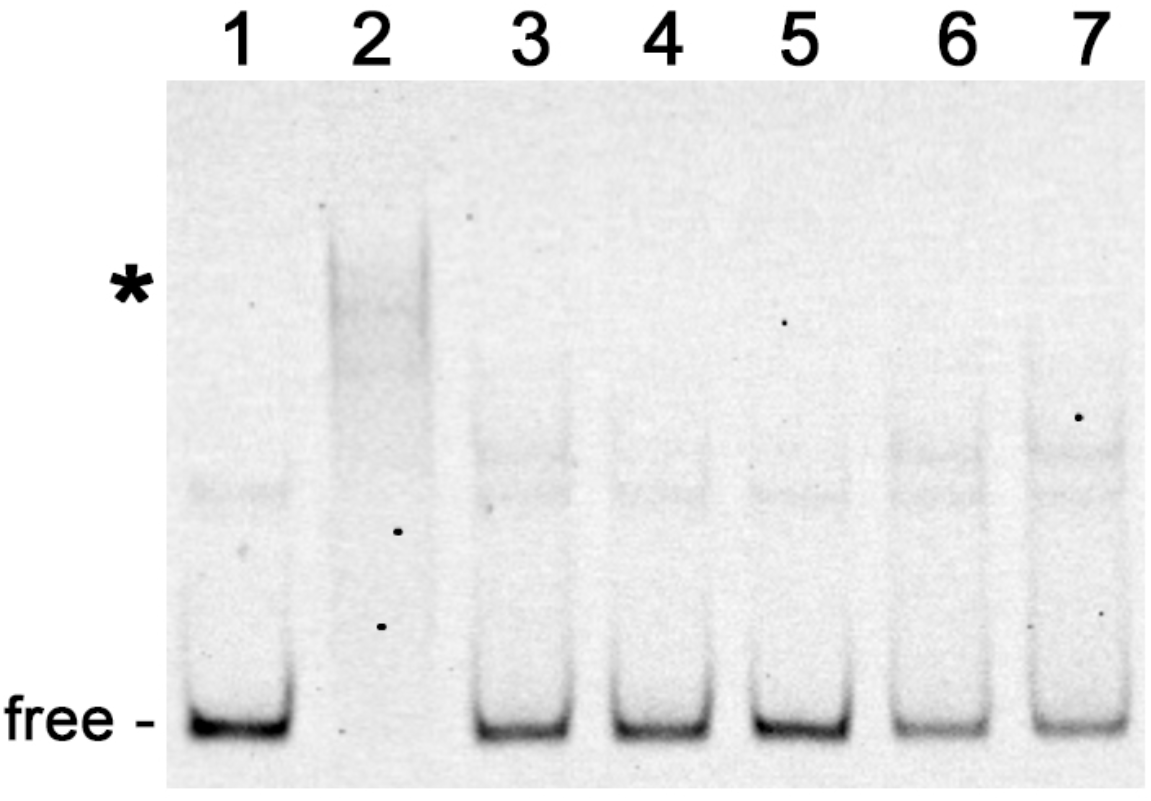
Representative EMSA of purified recombinant Gac with labeled B31 *ospC* operator/promoter DNA and unlabeled competitors. Shifted DNAs are indicated with asterisks. Unoccupied DNAs are labeled “free”. Labeled B31 operator/promoter (10 nM) without (lane 1) and with 25 nM Gac (lanes 2-7). Lane 2: no added competitor. Lane 3: plus 10x excess unlabeled B31 *ospC* operator/promoter. Lane 4: plus 25x excess unlabeled B31 *ospC* operator/promoter. Lane 5: plus 25x unlabeled pCR2.1 amplicon. Lane 6: plus 25x excess unlabeled *flaB* DNA. Lane 7: plus 25x unlabeled *ospAB* DNA.

**Supplemental Figure S2.**
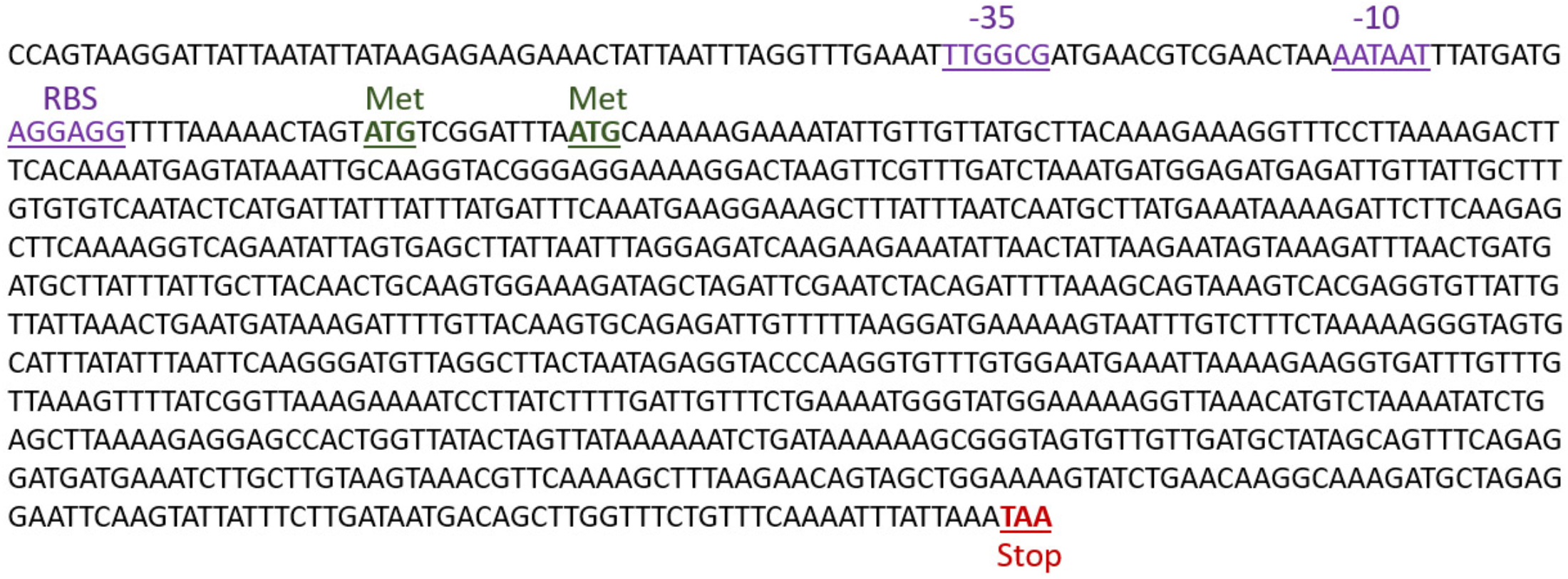
Sequence of the insert of pBLS820 that encodes Gac protein. Putative -10 and -35 promoter elements and the ribosome binding site (RBS) are indicated in **<u>purple</u>**. The *gac* ORF is preceded by two in-frame methionine codons, indicated in **<u>green</u>**. The *gac* termination codon is indicated in **<u>red</u>**.

